# DASC, a sensitive classifier for measuring discrete early stages in clathrin-mediated endocytosis

**DOI:** 10.1101/2020.01.28.924019

**Authors:** Xinxin Wang, Zhiming Chen, Marcel Mettlen, Jungsik Noh, Sandra L. Schmid, Gaudenz Danuser

## Abstract

Clathrin-mediated endocytosis (CME) in mammalian cells is driven by resilient machinery that includes >70 endocytic accessory proteins (EAP). Accordingly, perturbation of individual EAPs often results in minor effects on biochemical measurements of CME, thus providing inconclusive/misleading information regarding EAP function. Live-cell imaging can detect earlier roles of EAPs preceding cargo internalization; however, this approach has been limited because unambiguously distinguishing abortive-clathrin coats (ACs) from *bona fide* clathrin-coated pits (CCPs) is required but unaccomplished. Here, we develop a thermodynamics-inspired method, “disassembly asymmetry score classification (DASC)”, that unambiguously separates ACs from CCPs without an additional marker. After extensive verification, we use DASC-resolved ACs and CCPs to quantify CME progression in 11 EAP knockdown conditions. We show that DASC is a sensitive detector of phenotypic variation in CCP dynamics that is orthogonal to the variation in biochemical measurements of CME. Thus, DASC is an essential tool for uncovering the function of individual EAPs.

## Introduction

Clathrin-mediated endocytosis (CME) is the major pathway for cellular uptake of macro-molecular cargo (1). It is accomplished by concentrating cell surface receptors into specialized 100-200 nm wide patches at the plasma membrane created by a scaffold of assembled clathrin triskelia (2). The initiation and stabilization of these clathrin-coated pits (CCPs) is regulated by the AP2 (adaptor protein) complex (3), which recruits clathrin and binds to cargo and phosphatidylinositol-4,5-bisphosphate (PIP2) lipids. Numerous endocytic accessory proteins (EAPs), which modulate various aspects of CCP assembly and maturation, contribute to the formation of clathrin-coated vesicles (CCVs) that transport cargo to the cell interior. However, the exact functions of many of these EAPs are still poorly understood, and in some cases controversial (4, 5). Due to the resilience of CME, perturbing single EAPs, like CALM (6, 7), SNX9 (8, 9), etc. or even multiple EAPs (10) often results in minor/uninterpretable changes in bulk biochemical measurements of cargo uptake. Nonetheless, perturbed EAP functions can be physiologically consequential, *e.g*. CALM is identified as associated to Alzheimer’s disease (11) and SNX9 is correlated to cancer and other human diseases (12). We hence question whether measuring internalization by biochemical assays is sufficient for determining the actual phenotypes of missing EAP functions, and thereby further supporting clinical studies of the EAPs in more complex models.

Unlike bulk cargo uptake assays, the entire process of clathrin assembly at the plasma membrane can be monitored *in situ* by highly sensitive total internal reflection fluorescence microscopy (TIRFM) of cells expressing fiduciary markers for CCPs, such as the clathrin light chain fused to eGFP (13). Using this imaging approach, we and others have found that a large fraction of detected clathrin-coated structures (CSs) are shorter-lived (i.e. lifetimes < 20s) than thought to be required for loading and internalizing cargo, and dimmer (i.e. exhibit lower intensities) than mature CCPs detected prior to internalization (14, 15). These so-called “abortive” coats (ACs) presumably reflect variable success rates of initiation, stabilization and maturation, i.e. the critical early stages of CME. However, the range of lifetimes and intensities of ACs overlaps substantially with the range of lifetimes and intensities of productive CCPs (Fig. 1A,B). The current inability to unambiguously resolve ACs and CCPs limits analyses of the mechanisms governing CCP dynamics and their progression during CME.

**Figure 1.**
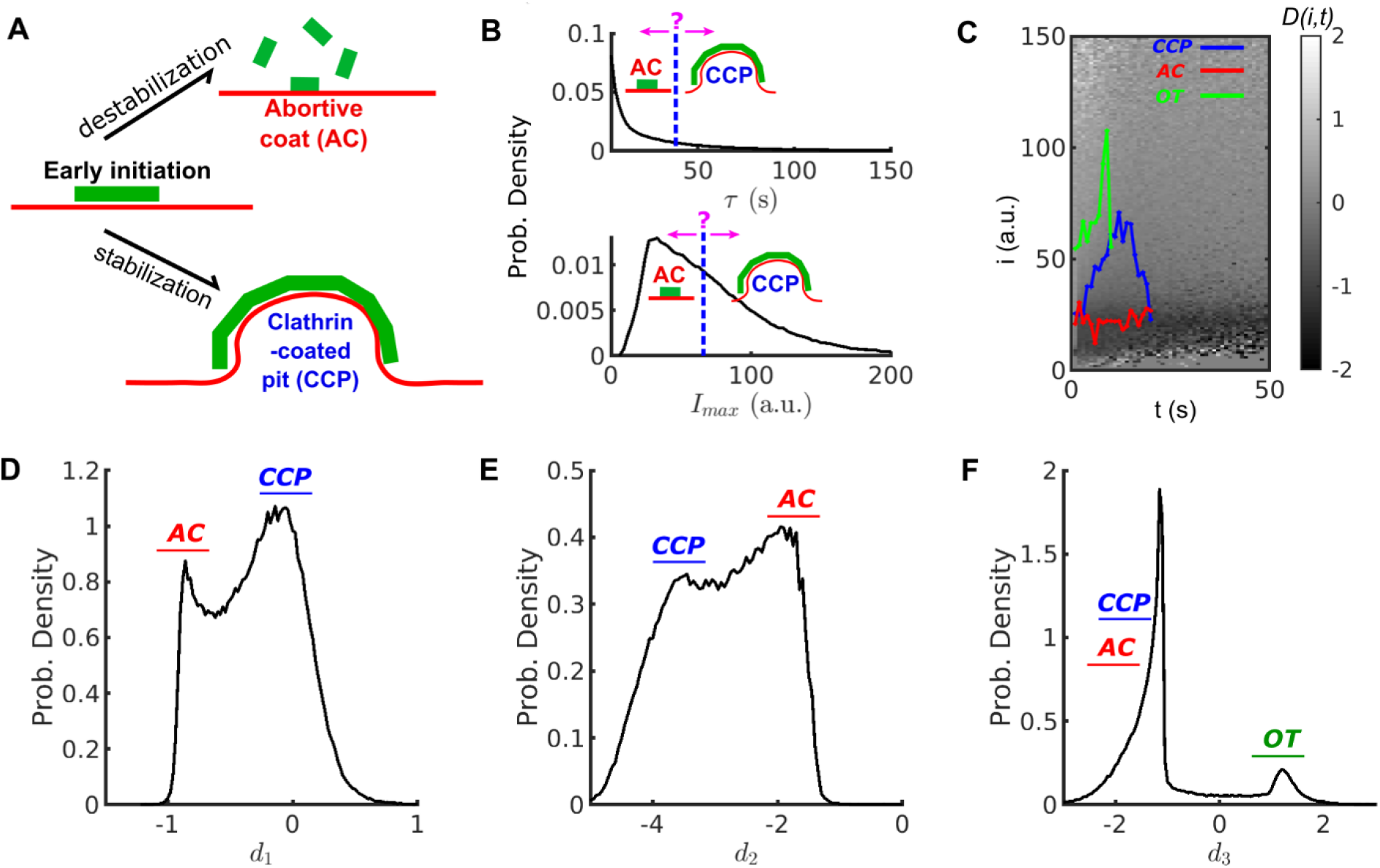
Conventional threshold based cut-off vs. DAS derived metrics. (A) Schematic of abortive coat (AC) and clathrin-coated pit (CCP) evolving from early clathrin nucleation. (B) Lifetime (*τ*) and intensity maxima (*I_max_*) characteristics of hypothetical ACs and CCPs. ACs are typically assigned by a user-defined lifetime or *I_max_* threshold. (C) Disassembly risk map *D*(*i*, *t*) represented on a gray value scale indicated by the gradient bar. A representative CCP (blue), AC (red) and outlier trace (OT) (green) are plotted on the *D*-map. (D) Distribution of *N* = 215,948 counts of *d*_1_ values for WT condition. AC group near *d*_1_ < 0 as a subpopulation, and CCP group at *d*_1_ ≈ 0 as another subpopulation. (E) Distribution of *N* counts of *d*_2_ values. Subpopulations of ACs and CCPs present in two modes. (F) Distribution of *N* counts of *d*_3_ values resolves the small subpopulation of OTs.

Our initial attempts to solve this problem relied on a statistical approach to deconvolve the overall broad lifetime distribution of all detected CSs into subpopulations with distinct lifetime modes (16). Although these statistical approaches allowed the identification of three kinetically-distinct CS subpopulations (16), the lifetimes of the thus identified subpopulations strongly overlapped, and the CS population with the longest average lifetimes, most likely representing productive CCPs, also contained a large fraction of very short-lived CCPs, which is structurally nonsensical. Later, as a result of improvements in the sensitivity of detection and tracking, eGFP-CLCa-labeled CSs were classified by imposing both lifetime and intensity thresholds (10, 17). Besides the subjectivity in setting these critical values, we demonstrate in this work that neither lifetime nor intensity are sufficient to classify CSs. More recently, Hong et al. (18) removed some subjectivity by training a Support Vector Machine (SVM)-based classifier of “false” vs “authentic” CCPs; but the underlying features were still largely based on lifetime and intensity thresholds, which themselves are sensitive to detection and tracking artefacts (see (10)). Finally, other efforts to distinguish abortive from productive events have introduced second markers, such as a late burst of dynamin recruitment (19, 20) or the internalization of pH sensitive-cargo (14) as classifiers, with the obvious drawbacks of more complicated experimental set ups. Clearly, the mechanistic analysis of CCP dynamics would greatly benefit from an objective and unbiased means to resolve these heterogeneous subpopulations.

Here, we introduce a thermodynamics-inspired method, referred to as *disassembly asymmetry score classification* (DASC), that resolves ACs from CCPs relying on the differential asymmetry in frame-by-frame intensity changes between disassembling and fluctuating/growing structures. DASC is independent of user-defined thresholds and prior assumptions, and does not require second markers. We confirmed the positive correlation between CCP stabilization and curvature generation by combining DASC with quantitative live cell TIRF and epifluorescence microscopy. We further applied DASC to phenotype siRNA-mediated knockdown of eleven reportedly early-acting ‘pioneer’ EAPs on CCP initiation and stabilization and compared these effects on CS dynamics with the effects on cargo uptake measured biochemically. In most cases we detected significant effects on early stages of CME resulting from reduced CCP initiation and/or stabilization that did not correlate with changes in transferrin uptake. Thus, DASC provides an orthogonal approach to traditional bulk biochemical measurements, and reveals compensatory mechanisms that can uncouple early perturbation from the final outcome of CME. Together these studies establish DASC as a new tool that is unique for objectively distinguishing abortive coats from *bona fide* CCPs and thus indispensable for comprehensively revealing which EAPs act at specific stages to mediate endocytic coated vesicle formation.

## Results

### Disassembly Asymmetry Score Classification (DASC): a new method to analyze CCP growth and stabilization

To ensure high sensitivity detection of all CCP initiation events, ARPE19/HPV16 (hereafter called HPV-RPE) cells were infected with lentivirus encoding an eGFP-tagged clathrin light chain a (eGFP-CLCa) and then selected for those that stably expressed eGFP-CLCa at ~5-fold over endogenous levels. Overexpression of eGFP-CLCa ensures near stoichiometric incorporation of fluorescently-labeled CLC into clathrin triskelia by displacing both endogenous CLCa and CLCb. Control experiments by numerous labs have established that under these conditions CME is unperturbed and that eGFP-CLCa serves as a robust fiduciary marker for coated pit dynamics at the plasma membrane (3, 10, 14, 16, 20–22). For all conditions, ≥19 independent movies were collected and the eGFP intensities of >200,000 clathrin structures per condition were tracked over time using TIRFM and established automated image analysis pipelines (10, 23). We refer to these time dependent intensities as traces. Each trace is a measure of the initiation, growth and maturation of the underlying clathrin structure (CS).

Following their initiation, the dynamics of CSs are highly heterogeneous, reflected by the widely spread distributions of lifetime and intensity maxima of their traces (Fig. 1B, top and bottom panels, respectively). Previous studies (10, 16, 20) have suggested that this heterogeneity reflects a mixture of at least two types of structures: 1) stabilized, *bona fide* CCPs, and 2) unstable partial and/or abortive coats (ACs) that rapidly turnover.

Productive CCPs (i.e. those that form CCVs and take up cargo) tend to have lifetimes >20s and reach an intensity level corresponding to a fully assembled coat (between 36 and >60 triskelia) (20). In contrast, ACs tend to exhibit lower intensity levels and disassemble at any time. However, CCPs and ACs strongly overlap in their lifetime and intensity distributions, especially during the critical first 20-30s after initiation. Consequently, the contributions of these two functionally distinct subpopulations of CSs to the overall lifetime or intensity distributions cannot be resolved and ACs cannot be readily distinguished from CCPs by application of a lifetime or intensity threshold (Fig. 1B). Significantly compounding the ability to distinguish CCPs from ACs is the fact that the intensities of individual CSs are highly fluctuating (see for example, Fig. S1A-B). These fluctuations are inevitable and reflect a combination of rapid turnover of individual triskelia, which occurs on the time scale of 1s (1, 24), stochastic bleaching of fluorophores, and membrane fluctuations within the TIRF field. We thus sought an approach to distinguish ACs from *bona fide* CCPs that is independent of user-defined thresholds and leverages these intensity fluctuations measured at high temporal resolution.

Inspired by the computation of entropy production (EP) (25) we designed a new metric derived from the fluctuations of clathrin intensity traces that can clearly separate ACs from CCPs. Conventionally, EP quantifies the dissipation rate of thermal energy when a system of interest is driven far away from equilibrium, as is the case during the formation of a macro-molecular assembly such as a CCP. This quantity is obtained by computing the difference between forward and reverse reaction rates. We therefore assigned clathrin assembly and disassembly as forward and reverse reactions in order to derive an EP-based metric of the progression of CS formation.

We first expressed each trace as a chain of transitions among integer intensities (or states) over time, for the *n*th trace,

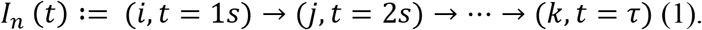

In this example, *i*, *j*… *k* ∈ [1, *i_max_*](*a*. *u*.), where *i_max_* is the largest intensity recorded so that [1, *i_max_*] represents the entire pool of the intensity states. *τ* is the lifetime of this trace (see Materials and Methods for details).

Next, after expressing all the traces as in eq. (1), we quantified for each transition between two intensity states the conditional probabilities *W_t_*(*i*^⊖^|*i*) and *W_t_*(*i*|*i*^⊖^). Given state *i* and its ***lower states i***^⊖^ ∈ [1, *i* − 1], *W_t_*(*i*^⊖^|*i*) denotes the probability of a decrease in intensity *i* → *i*^⊖^ between time *t* to *t* + 1, and *W_t_*(*i*|*i*^⊖^) denotes the probability for an increase in intensity *i*^⊖^ → *i* (see Materials and Methods for details).

From these probabilities, we define a disassembly risk function (*D*) for any given intensity-time state (*i*, *t*) as:

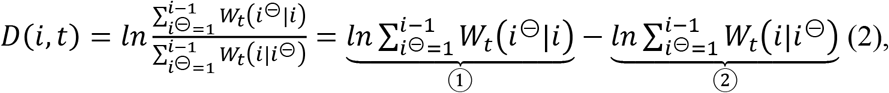

where, between state *i* and its lower states *i*^⊖^ at time *t*, Term ① includes every transition of *clathrin loss*; while Term ② includes every transition of *clathrin gain*. *D* = ① – ② thus indicates the net risk for disassembly at every intensity-time state.

We can use this *D* function to project each trace into a space of disassembly risk (Fig. 1C). The projected trace (Fig. S1C) then predicts the disassembly risk for an individual trace of particular intensity at a specific time. For example, *I_n_* (*t*) in eq. (1) yields a corresponding series of disassembly risk (see Fig. S1C), written as:

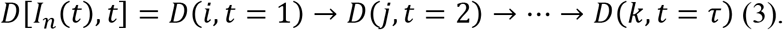

Hence, each intensity trace as in eq. (1) is translated into a *D* series reflecting the risk of disassembly at each time point. Most *D*(*i*, *t*) values are either negative (low disassembly risk, i.e. loss < gain) or nearly zero (moderate disassembly risk), see Fig. 1C, which we interpret as reflective of two phases of CCP growth and maturation.

1. **Early growth phase:** Following an initiation event, and during the first few seconds of CME, almost all CSs, including ACs, grow albeit with fluctuation. Also, most CSs are still small. Hence, in this earliest phase, Term ① < Term ② and *D*(*i*, *t*) < 0. Accordingly, clathrin dissociation is rare and all traces in this early phase have a low risk of disassembly. However, the risk of acute disassembly increases as CSs approach the end of this phase. CSs that disassemble early are potentially ACs, whereas surviving CSs enter the next phase to become CCPs.
2. **Maturation phase:** Upon completion of the growth phase, CCP intensities plateau but continue to fluctuate over many high intensity states at mid to late time points. The fluctuation is equivalent to having a similar chance of gain or loss of clathrin, thus Term ① ≈ ② and *D*(*i*, *t*) ≈ 0. CCPs in this phase retain a moderate risk of acute disassembly.

In summary, *D*(*i*, *t*) < 0 is indicative of early stages of clathrin recruitment when disassembly risk is suppressed; *D*(*i*, *t*) = 0 is indicative of intensity fluctuations that occur at later stages of CCP growth and maturation. Fig. 1C displays representative examples of CCP (blue) and AC (red) traces. In the early growth phase both traces exhibit *D*(*i*, *t*) < 0 (dark shaded background). As CCPs reach the maturation phase they approach the regime *D*(*i*, *t*) ≈ 0. Thus, even short-lived CCPs tend to have larger *D* values than ACs and for longer-lived CCPs the contribution of the early growth phase with negative *D* values becomes negligible. Accordingly, for maturing CCPs *D* values distribute around zero, whereas for ACs *D* values distribute in the negative range.

A small portion of CSs possess abnormally high intensities when first detected, but quickly disappear. Therefore, Term ① > Term ②, *D*(*i*, *t*) > 0, and the disassembly risk for high intensity states at early time points is high (green traces in Fig. 1C and Fig. S1C). These atypical CSs frequently appear in regions of high background, which can obscure early and late detections (Fig. S1D) and impair the ability to accurately detect small intensity fluctuations. As interpreting the fates of these CSs is difficult, and because they are rare, we refer to them as **outlier traces** (OTs).

To quantitatively distinguish the distributions of CCPs and ACs, we examined mean, variation and skewness of the *D* series. Considering the *n*th series *D*[*I_n_*(*t*), *t*], we first calculated its time average:

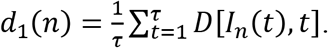

An AC is expected to have *d*_1_(*n*) < 0, whereas a CCP is expected to have *d*_1_(*n*) ≈ 0. Indeed, for a population of *N* > 200,000 CSs tracked in HPV-RPE cells, the distribution of *d*_1_ values is bimodal (Fig. 1D), allowing the distinction of ACs and CCPs.

We additionally computed:

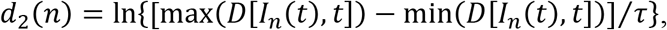

which reflects the lifetime-normalized difference between the maximum and minimum value of a *D* series. For example, the *D*-series of the CCP trace in Fig. 1C (see blue curve in Fig. S1C) has a maximum value of 0.2 and minimum value of −0.8, and lasts for 30s (Fig. S1C). Thus 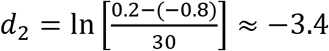. Analogously, the *D*-series of the AC trace (red curve in Fig. S1C) yields 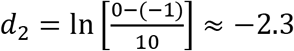. In general, because traces of ACs are dominated by the early growth phase with *D* continuously changing, they are expected to have a significantly greater *d*_2_ value than CCPs. Indeed, the distribution of this feature is also bimodal (Fig. 1E) and thus can strengthen the distinction between ACs and CCPs.

The *D* series of OTs contain a few initial values that are much higher than those in the *D* series associated with either ACs or CCPs (Fig. S1C). Therefore, such series can be identified via a modified skewness of *D*:

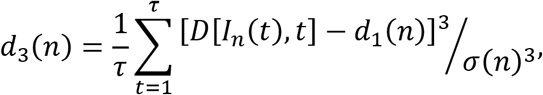

where 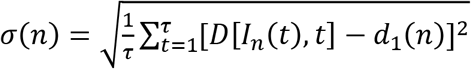 is the standard deviation of the *D* series.

Indeed, the distribution of *d*_3_ over *N* series displays two tight populations with the *d*_3_ values of OTs easily separable from the *d*_3_ values of ACs and CCPs (Fig. 1F).

Using the three summary statistics (*d*_1_, *d*_2_, *d*_3_) we project all CS traces into a feature space (Fig. 2A) and classify ACs (red), CCPs (blue), and OTs (green) using k-medoid clustering (see Materials and Methods). Values for *d*_3_ identify OTs, whereas *d*_1_ and *d*_2_ complement one another separating ACs from CCPs. As these features originate from the disproportionate disassembly vs. assembly of CSs, we term our feature selection the *disassembly asymmetry score* (DAS), and name the packaged software DASC as DAS classification, available under https://github.com/DanuserLab/cmeAnalysis.

**Figure 2.**
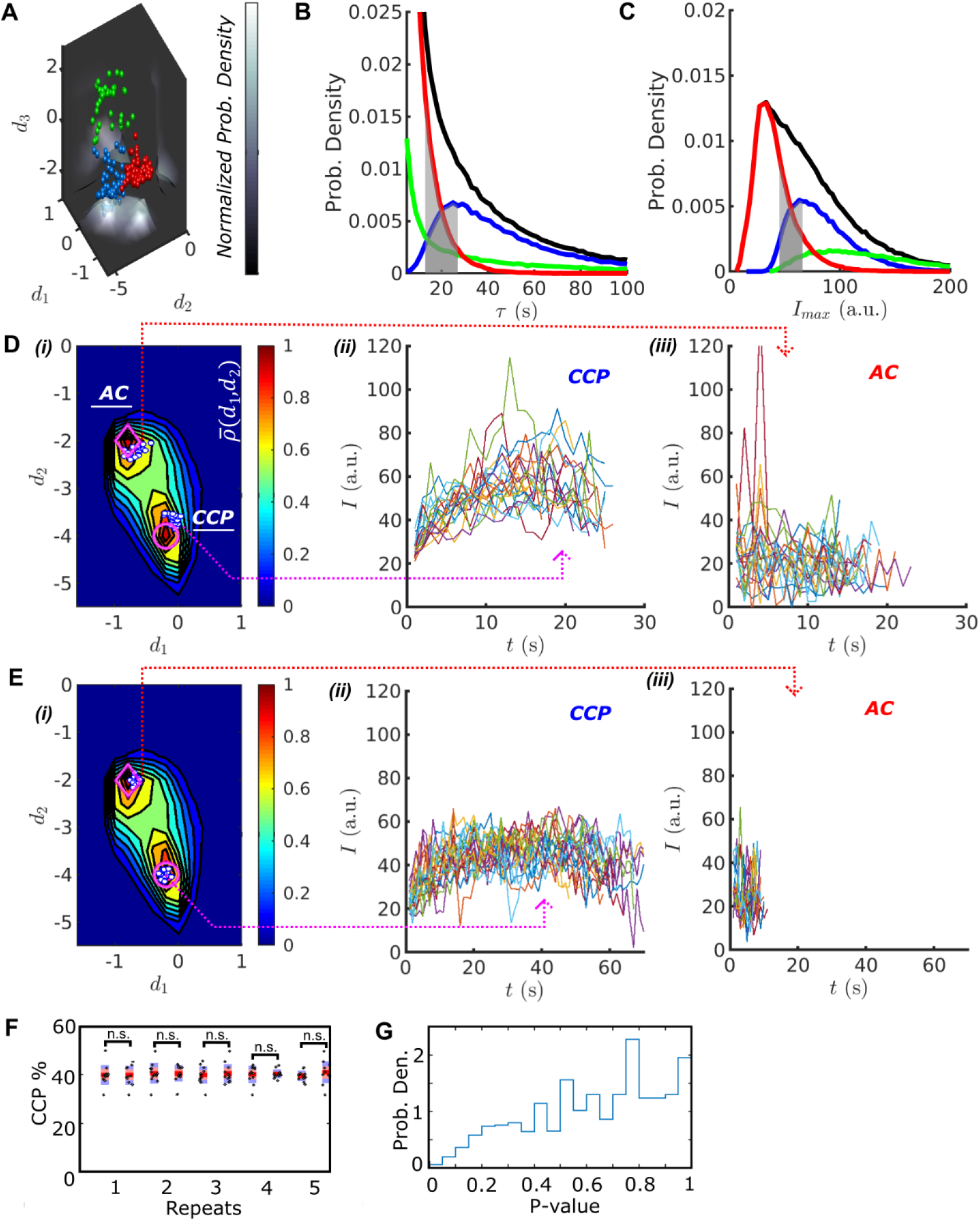
DASC resolves behaviorally distinct ACs and CCPs. (A) k-medoid classification in three-dimensional feature space (***d_1_***, ***d_2_***, ***d_3_***), where normalized probability densities 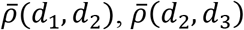 and 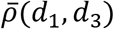 are shown as three landscape plots. 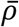 values are scaled according to gray bar. Examples of CCPs (blue), ACs (red) and OTs (green) concentrate near the maxima of 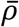. (B) Lifetime distributions of all traces (black), CCPs (blue), ACs (red) and OTs (green). Gray region shows lifetime overlap between CCPs and ACs. (C) *I_max_* distributions and overlap. Color scheme same as in (B). Overlap between CCPs and ACs. (D) i. DAS plot: 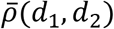 contour map (values indicated by color bar) with modes for CCPs and ACs indicated by circle and diamond, respectively. Ten representative CCPs and ACs (white dots) from the lifetime overlap in (B) close to the modes are projected onto *d*_1_-*d*_2_ coordinate. Traces of the representative CCPs (ii) and ACs (iii) from i. (E) Same as (D) for the representative CCPs and ACs from the *I_max_* overlap in (C). (F) Five examples of comparison between 12 and another 12 movies of WT cells imaged on the same day. A total of 24 movies were randomly shuffled to obtained 12-12 pairs. (G) p-value distribution of 1000 repeats of shuffle.

### DASC accurately identifies dynamically distinct CS subpopulations

The DASC-resolved subpopulations of CSs exhibit distinct but overlapping lifetimes and intensities (Fig. 2B, C grey zone), confirming the inability of these conventional metrics to distinguish ACs from CCPs. The lifetimes of ACs (Fig. 2B, red) were predominantly short (<20s) and exhibited an exponential distribution characteristic of coats that are exposed to an unregulated disassembly process. In contrast, CCP lifetimes (Fig. 2B, blue) exhibited a unimodal distribution with a highest probability lifetime of ~26s. In previous work, we had shown that this distribution is best represented by a Rayleigh distribution that reflects the kinetics of a three-to four-step maturation process (10, 16, 26). Interestingly, although partially overlapping with ACs, the intensity distribution of CCPs (Fig. 2C) exhibits a sharp threshold in the minimum intensity, indicative of the minimum number of clathrin triskelia required to form a complete clathrin basket (19, 20). The majority of OTs (Fig. 2B, C, green) are highly transient and bright structures, reflective of the higher backgrounds from which they emerge.

Despite their overlapping lifetimes and intensities, AC and CCP traces are well resolved by DASC as represented in two-dimensional, normalized probability density maps 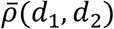 (Fig. 2Di), from here on referred to as DAS plots (see Materials and Methods for details). To illustrate this point, we selected 10 CSs with overlapping lifetime distributions (10-25s, gray zone Fig. 2B) that fall close to the associated modes of either the AC or CCP populations in the DAS plot, i.e. the two maxima of 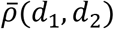 denoted by a diamond for ACs and circle for CCPs in Fig. 2Di. White dots show the (*d*_1_, *d*_2_) locations of the selected CSs. Their intensity traces are shown in Fig. 2 Dii-iii. Although the lifetimes are almost identical, the CCP and AC traces show characteristic differences in their intensity evolution. CCP intensities rise to a clear maximum as they assemble a complete clathrin coat (Fig. 2Dii), followed by a falling limb, which is associated with CCV internalization and/or uncoating. In contrast, AC intensities are more random (Fig. 2Diii), suggesting that these coats, trapped in the early growth phase, undergo continuous exchange of clathrin subunits without significant net assembly. We occasionally observed rapidly fluctuating, high intensity CSs amongst the AC traces. These likely correspond to previously identified ‘visitors’ (i.e. endosome-associated coats transiting through the TIRF field), which make up ~10% of all detected CSs (10).

We next selected 10 CSs (indicated as white dots in Fig. 2Ei) from the AC and CCP populations that fall into the overlap region in the distributions of intensity maxima (gray zone, Fig. 2C). Although the intensity ranges are nearly identical, the selected CCP traces again display a rising and falling limb and lifetimes of ~60s (Fig. 2Eii). In sharp contrast, ACs fluctuate about the same intensity values (Fig. 2Eiii) and exhibit much shorter lifetimes of ~10s. Together, these data demonstrate that DAS provides an unbiased metric to discriminate between two completely distinct clathrin coat assembly and disassembly processes that, by inference, are associated with abortive coats and *bona fide* CCPs. This has not been possible based on more conventional features such as lifetime and intensity (10, 16–18, 20, 27).

To further establish the robustness of the DASC, we acquired 24 movies from the same WT condition on the same day and randomly separated them into pairs of 12 movies each. We then applied DASC to the movies, and calculated percent contribution of *bona fide* CCPs, i.e. CCP% = CCP:(CCP+AC+OT) × 100%, for each movie using the first 12 as the control set, the other 12 as the test set, and compared the two data sets. The procedure was repeated 1000 times. Fig. 2F shows 5 example pairs, and Fig. 2G gives the p-value distribution of the 1000 repeats. As expected, comparison by a Wilcoxon rank sum test yields p-values >0.5 for most pairs and scarcely <0.1, indicating no significant difference between movies from the same condition. Thus, DASC is statistically robust and not overly sensitive to movie-to-movie variations in data collected on the same day.

### Validation through perturbation of established CCP initiation and stabilization pathways

We next tested the performance of DASC against conditions known to perturb early stages in CME. AP2 complexes recruit clathrin to the plasma membrane and undergo a series of allosterically-regulated conformational changes needed to stabilize nascent CCPs (17, 28–30). Previous studies based on siRNA-mediated knockdown (KD) of the α subunit of AP2 and reconstitution with either WT, designated αAP2(WT), or a mutant defective in PIP2 binding, designated αAP2(PIP2^-^), in hTERT-RPE cells have established that αAP2-PIP2 interactions are critical mediators of AP2 activity (17). We repeated these experiments in HPV-RPE cells using DASC and detected pronounced differences in the DAS plots derived from αAP2(WT) vs αAP2(PIP2^-^) cells (Fig. 3Ai-ii). A DAS difference Δ*ρ*[*αAP2*(*WT*), *αAP2*(*PIP2*^−^)] map (see Materials and Methods) shows a dramatic increase (yellow) in the fraction of ACs and a corresponding decrease (black) in the fraction of CCPs (Fig. 3B), as expected given the known role of αAP2-PIP2 interactions in CCP stabilization.

**Figure 3.**
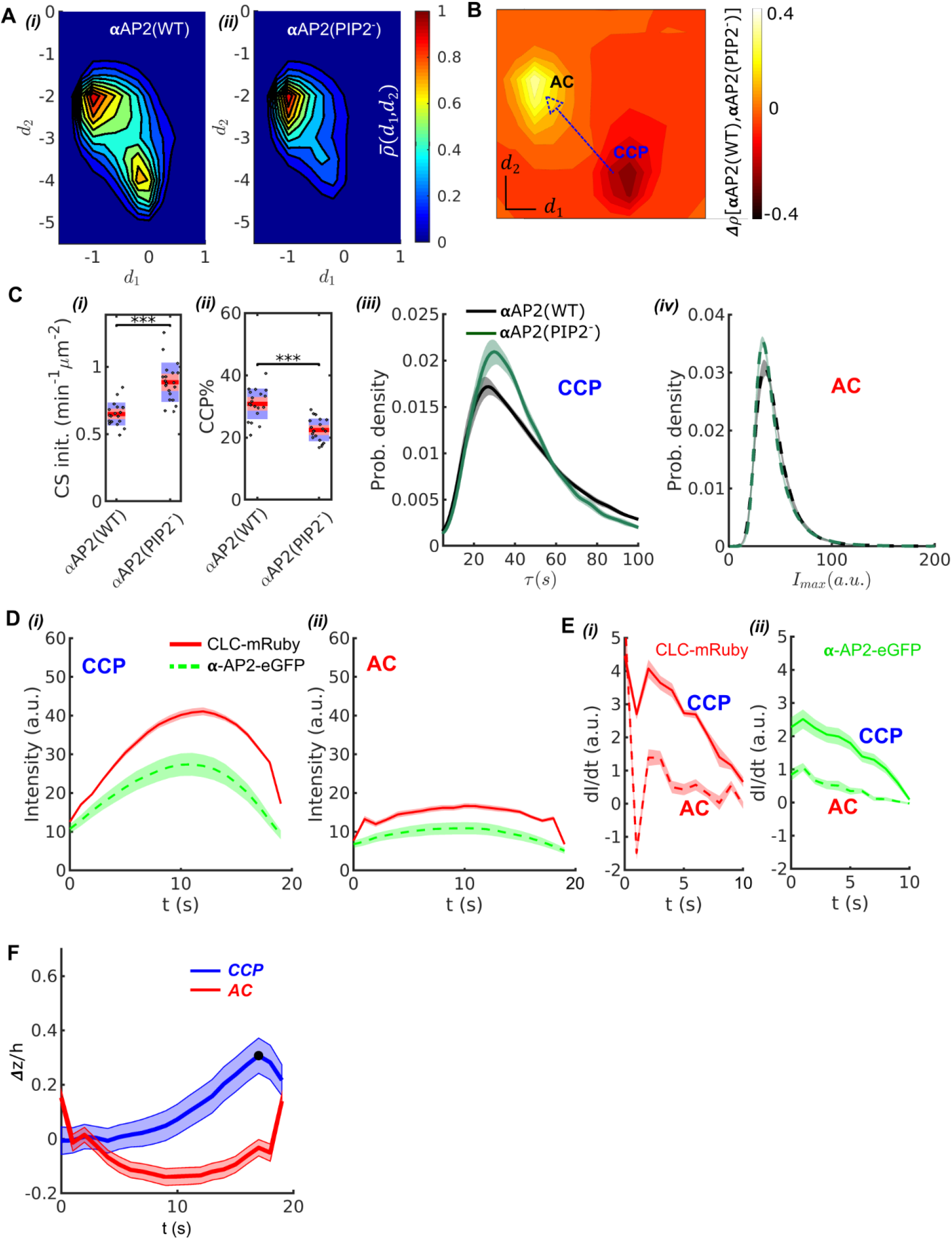
Validation of DASC. (A) DAS plots showing 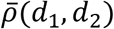 contour as ‘rainbow’ map and color bar for αAP2(WT) cells (i) and αAP2(PIP2-) cells (ii). (B) DAS difference plot (difference in *d*_1_*d*_2_ distribution) of αAP2(PIP2-) minus αAP2(WT) cells as contour in ‘heat’ map. (C) Comparison of DASC-derived metrics for CCP dynamics in αAP2(WT) vs αAP2(PIP2-) cells. CS initiation rate (i) and CCP% (ii), population ratio as percentage: [*n_CCP_*/(*n_CCP_* + *n_AC_* + *n_OT_*)] × 100 in αAP2(WT) and αAP2(PIP2-) cells. Dots represent jittered raw data from individual movies, box plots show mean as red line and 95% and 1 standard deviation as red and blue blocks, respectively (see Materials and Methods). (iii) CCP lifetime distribution of αAP2(WT) vs αAP2(PIP2-) cells. (iv) *I_max_* distribution of ACs in αAP2(WT) vs αAP2(PIP2-) cells. (D) 20 second cohorts from dual channel movies of CLC-mRuby (red, solid) and α-AP2-eGFP (green, dashed) for CCPs (i) and ACs (ii). (E) Time derivative of CLC-mRuby (i) and α-AP2-eGFP (ii) intensities for the first 10 seconds in the dual channel cohorts of CCPs and ACs in (D). (F) Time course of invagination depth Δ*z*(*t*)/*h* (TIRF characteristic depth *h* = 115*nm*) for CCPs (blue) and ACs (red) measured by Epi-TIRF. Statistical analysis of the data used the Wilcoxon rank sum test. *** p-value < 0.001, ** p-value < 0.01, * p-value < 0.05, n.s. (non-significant) p-value > 0.05. Shaded area indicates 95% confidence interval.

We also observed an increase in CS initiation rate (CS init.) (Fig. 3Ci), measured by total trackable CSs detected per minute per cell surface area (see Material and Methods for detail definition). Previous studies reported a decrease in CS initiation rate (17). This apparent discrepancy likely reflects our use of all detected traces to calculate CS initiation rate as compared to previous use of only ‘valid’ tracks (see Materials and Methods). As the CSs observed in the αAP2-PIP2^-^ cells were significantly dimmer than those detected in WT cells (see Fig. S2), more initiation events would have been scored as ‘invalid’ in the previous analysis due to flawed detections, especially at early stages of CCP assembly.

DASC analysis revealed multiple defects in early stages of CME in the αAP2(PIP2^-^) cells compared to αAP2(WT) cells. We detected a pronounced decrease in the efficiency of CCP stabilization, which was calculated as the fraction of CCPs in all the valid traces (see Material and Methods). The CCP% decreased from 32% (on average) in control cells to 22% in αAP2(PIP2^-^) cells (Fig. 3Cii). The lifetime distributions of CCPs also shift to shorter lifetimes (Fig. 3Ciii), resulting in decreased median lifetimes (Table 1) in αAP2(PIP2^-^) cells compared to αAP2(WT) cells. This lifetime shift indicates that the mutation can also cause instability in fully grown clathrin coats, as previously reported (17). In addition, the maximal intensity of ACs, which is an indication of the growth of ACs before they are turned over, was reduced in αAP2(PIP2^-^) cells (Fig. 3Civ, Table 1). These data suggest greater instability of nascent coats.

**Table 1.** Quantitative summary of EAP experiments. ↑ = increase; ↓ = decrease; → = no significant change, p-value>.05; *** p-value<.001; ** p-value<.01; * p-value<.05 (statistical tests explained in Materials and Methods), percentage change based on mean-mean comparison between KD and control condition.

| EAP | Number of movies | Initiation | Stabilization | Initiation + Stabilization | Maturation | Biochemical measurements of CME |  |
| --- | --- | --- | --- | --- | --- | --- | --- |
| siRNA or mutant | $n_{siCtrl}$<br>$n_{siEAP}$ | CS initiation rate | CCP% | CCP rate | Median lifetime of CCP | *TfRint* (intracellular accumulation) | *TfReff* (internal/surface bound) |
| α-PIP2 | 19, 20 | ↑36%*** | ↓27%*** | ↑27%** | ↓* | -- | -- |
| CALM | 20, 19 | ↓28%*** | ↓34%*** | ↓62%*** | ↑** | ↑21%*** | ↓64%*** |
| epsin1 | 23, 22 | → | ↓27%*** | ↓27%*** | → | ↓37%*** | → |
| Eps15 | 23, 23 | ↓25%*** | ↓29%*** | ↓48%*** | → | → | ↑22%** |
| Eps15R | 23, 24 | ↓24%*** | ↓14%*** | ↓37%*** | → | ↓13%*** | → |
| FCHO1 | 24, 24 | ↑15%** | → | ↑12%* | → | ↓30%*** | → |
| FCHO2 | 24, 24 | → | ↓19%*** | → | → | ↓22%*** | ↓34%*** |
| ITSN1 | 20, 19 | → | ↓26%*** | ↓31%** | → | ↓22%*** | ↓9%* |
| ITSN2 | 20, 22 | ↓13%** | ↓8%* | ↓37%*** | ↑* | ↓31%*** | ↓21%*** |
| NECAP1 | 22, 21 | ↓13%*** | ↓23%*** | ↓34%*** | ↑*** | → | ↓24%** |
| NECAP2 | 22, 21 | → | → | → | → | → | ↓13%** |
| SNX9 | 24, 24 | ↓31%*** | ↓45%*** | ↓67%*** | ↑*** | ↑21%*** | ↓57%*** |

We next compared the kinetics and extent of recruitment of AP2 and clathrin to ACs and CCPs. For this we applied two-color imaging and ‘master-slave’ analysis (10) to simultaneously track clathrin and AP2 in ARPE cells stably expressing mRuby2-CLC as the master channel and the wild-type α subunit of AP2 encoding eGFP within its flexible linker region (α-eGFP-AP2) as the slave channel. Applying DASC to the mRuby2-CLCa signal to distinguish CCPs from ACs and cohort plotting of traces with lifetimes in the range 15s to 25s (26), we observed, as expected, that CCPs reach significantly higher average clathrin intensity than ACs (Fig. 3Di-ii). We also observed significantly higher levels of AP2 α subunit present at CCPs than ACs. Moreover, as previously shown for statistically-defined abortive vs. productive pits (26), the initial rates of recruitment to CSs of both clathrin and AP2, determined by the derivative of intensity, were much greater for CCPs than ACs (Fig. 3Ei-ii). Together these data corroborate the known stabilization function of AP2 during CCP initiation (17, 31), and serve to validate the ability of DASC to distinguish different regimes of molecular regulation at early stages of CME.

### Validation through curvature acquisition and CCP stabilization relation

Previous studies have suggested that curvature generation within nascent CCPs is a critical factor for their maturation and that CCPs that fail to gain curvature are aborted (10, 16, 27, 32). Therefore, we asked how DASC-identified ACs and CCPs relate to the acquisition of CCP curvature (Fig. 3F). To this end, we applied DASC to traces acquired by near simultaneous epifluorescence (Epi)-TIRF microscopy (10, 26). Because of the differential fluorescence excitation depths of TIRF- and epi-illumination fields, the ratio of Epi:TIRF intensities of individual CSs provides a measure of curvature (Fig. S3A). CSs were classified as ACs or CCPs based on the TIRF channel traces and then grouped into lifetime cohorts to obtain average invagination depth Δ*z* (See Methods and materials). We show in Fig. 3F that CSs in the 20s cohort classified by DASC as CCPs reached maxima Δ*z_max_*/*h* = max[Δ*z*(*t*)]/*h* > 0.3, which corresponds to an invagination depth of > 35*nm* (*h* = 115*nm* is the characteristic depth of our TIRF illumination field, see Materials and Methods). In contrast, CSs in 20s cohort classified by DASC as ACs fail to gain significant curvature. Other cohorts supporting this CCP-curvature relation are presented in Fig. S3. Together, these data (Fig. 3D, F) establish that DASC-identified ACs and CCPs are structurally and functionally distinct.

### Differential effects of endocytic accessory proteins (EAPs) on CCP dynamics revealed by DASC

Equipped with DASC as a robust and validated tool to distinguish *bona fide* CCPs from ACs and to quantitatively measure early stages of CME, we next probed the effects of siRNA KD of eleven EAPs previously implicated in these stages (3, 33–43). Our measurements allow us to segment the early dynamics in CME into discrete stages (Fig. 4A), including stage 1: initiation, measured by *CS initiation rate* (*CS init. in min^-1^ μm^-2^*), and stage 2: stabilization, quantified by *CCP*%, which is a measure of the efficiency of nascent CCP stabilization (Fig. 4A). Combining stage 1 and 2 measurements, we calculated *CCP rate* (*min^-1^ μm^-2^*), *i.e*. the number of CCPs appearing per unit time normalized by cell area (see Materials and Methods for a detailed definition and computation of the three metrics). Finally, we measured the lifetime distribution of CCPs, which reflects CCP maturation (stage 3) and the efficiency of transferrin receptor (*TfR*) uptake, *TfReff*, a bulk measurement of internalized *TfRs* as a percentage of their total surface levels, which is not stage specific but reflects the overall process of CME (see Fig. 4A, Fig. S4 and Materials and Methods).

**Figure 4.**
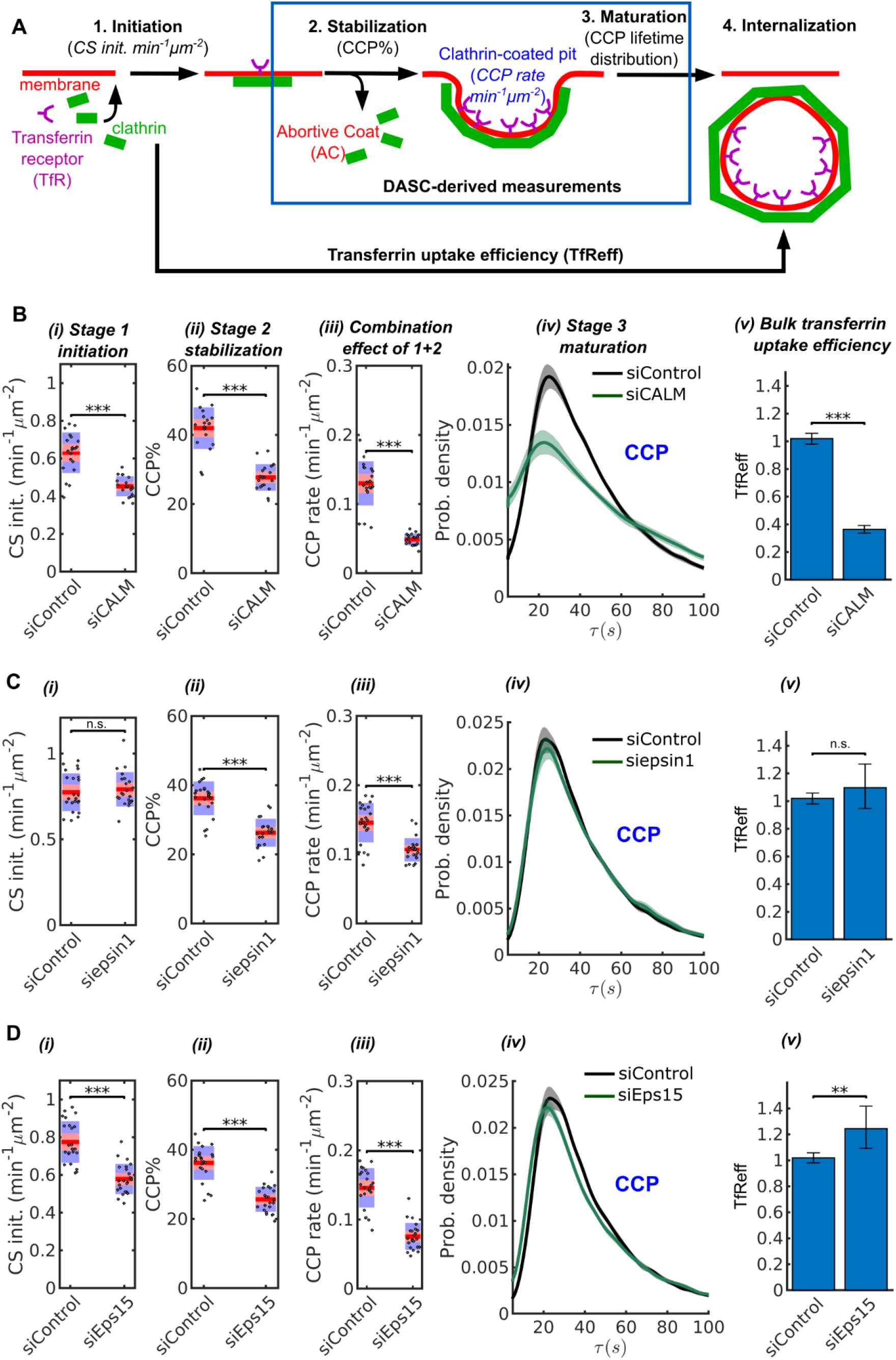
Stage specific phenotypes detected by DASC compared to transferrin uptake measurement. (A) Schematic of 4 stages of CME: CS initiation, CCP stabilization, CCP maturation and CCV internalization. Stage 1-3 are quantified by CS initiation rate (CS init. in *min^-1^ μm^-2^*), CCP% and CCP lifetime distribution. Bulk assays for transferrin receptor uptake (*TfReff*) measure CCV formation are not stage specific. CCP rate (*min^-1^ μm^-2^*) measures the combination of initiation and stabilization. Effects of siRNA knockdown of CALM (B), epsin1 (C) and Eps15 (D) on (i) CS initiation rate, (ii) CCP%, (iii) CCP rate, (iv) CCP lifetime distribution and (v) *TfReff* (internalized over surface bound transferrin receptors, error bars as 95% confidence interval and statistical significance explained in Materials and Methods).

The KD effects of these EAPs are summarized in Table 1 with statistical significance. KD efficiency is shown in Fig. S4. Three representative examples comparing the effects of treating cells with EAP-specific siRNA vs non-targeting siRNA on i) stage 1-initiation, ii) stage 2-stabilization, iii) stage 1 plus 2, iv) stage 3-maturation and v) transferrin uptake efficiency (*TfReff*) are shown in Fig4 B-D. KD of CALM dramatically decreased initiation, stabilization and *TfReff*, and also significantly altered the lifetime distribution of CCPs (Fig. 4B). These changes included increases in both short- and long-lived CCPs, indicative of a role for CALM in multiple aspects of CCP maturation. Conversely, KD of epsin1 selectively perturbed CCP stabilization without affecting initiation, CCP lifetime or *TfReff* (Fig. 4C). In the example of Eps15, initiation and stabilization were significantly decreased upon KD, while CCP lifetime was not significantly affected; on the other hand, *TfReff* was slightly increased (Fig. 4D), suggesting a compensation effect. Together, these examples show consistent and significant defects in early stage(s) caused by the three EAPs, despite their differential and less interpretable effects on the efficiency of transferrin receptor uptake.

Moreover, DASC analysis confirmed that KD of the so-called ‘pioneer’ EAPs, e.g. FCHO1/2, ITSN1/2, NECAP1 and Eps15/15R (34) selectively altered CCP initiation and/or stabilization without affecting CCP maturation rates, and having only relatively mild effects on the efficiency of *TfR* uptake (Table 1). In sum, DASC is a statistically reliable method to detect phenotypes caused by KD of individual EAPs, thus enabling their effects on specific stage(s) of CCP dynamics to be mechanistically dissected.

### DASC phenotypes are orthogonal to biochemical measurements of CME efficiency

We next evaluated the sensitivity of DASC and its relation to bulk biochemical measurement of transferrin uptake (*TfReff*), the commonly used assessment of CME efficiency. Strikingly, KD of most EAPs significantly reduced *CCP rate* (by over 30%), but caused less and/or uncorrelated shifts in *TfReff* (Fig. 4 panel (iii), (v) and Table 1). To further explore this observation, we first replotted the KD phenotypes as percentage changes relative to control (Δ_r_) in a colored ‘heat’ map (Fig. 5A). We also added measurements of transferrin receptor internalization (*TfRint*), which is independent of potential changes in surface levels of the recycling *TfR*, as is often measured by FACS or fluorescence imaging. As is evident from this plot, DASC-determined changes to early stages, Δ_r_*CCP*% and especially Δ_r_*CCP rate*, were with few exceptions more severe than Δ_r_ *TfRint* and Δ_r_*TfReff*. Few of the pioneer proteins we studied affected CCP median lifetime (Δ_r_τ_CCP_) and thus later stages of CCP maturation.

**Figure 5.**
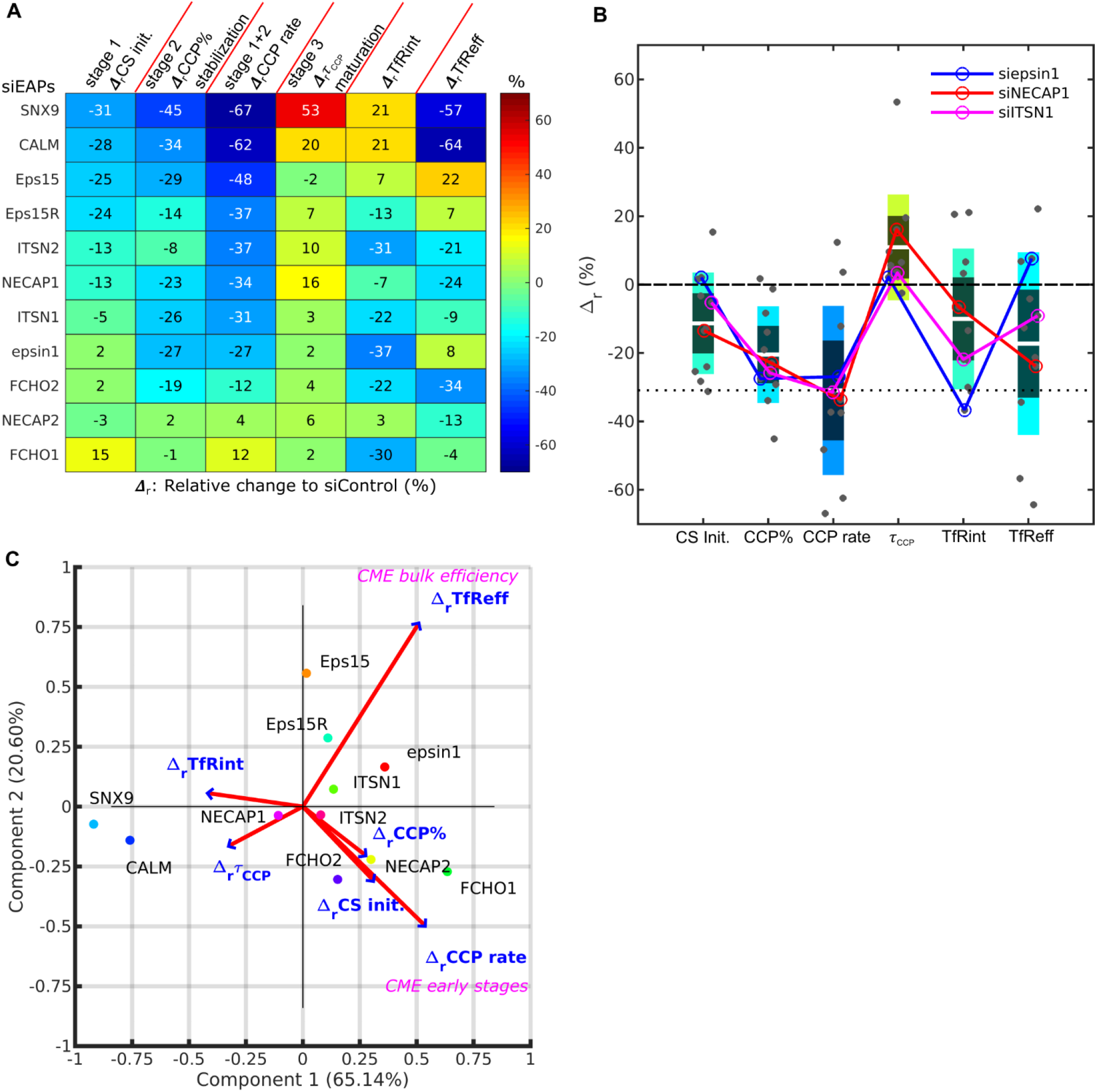
DASC is a sensitive measure of stage-specific defects in CME not detected by bulk measurement of transferrin uptake. (A) Summary of phenotypes of 11 EAP KD conditions evaluated by percentage difference (Δ_r_) in CS initiation rate (CS init.), CCP%, CCP rate, CCP median lifetime (τ_CCP_) and transferrin receptor uptake: internalized and efficiency (TfRint and TfReff) relative to control. EAP KD sorted based on CCP rate. (B) Bar graph of 6 variables of Δ_r_ in (A). Each bar colored based on its mean Δ_r_ value matching to color bar in (A). 3 example conditions, KD of epsin, NECAP1 and ITSN1, presented as circles plus lines. Δ_r_ = 0 presented as dashed line, averaged Δ_r_ CCP rate as dotted line. (C) Principle component analysis (PCA). Projection of 6 variable values from 11 conditions in (A) into principle component space. First and second component (Component 1 and 2) account for 65.14% and 20.60% of total variance, respectively. Projection of original variable axes presented as red vectors with blue arrows.

To visualize the range of effects of all EAPs on each measurement, we next plotted the data from Fig. 5A in a bar graph (Fig. 5B, each black dot represents an EAP KD). Three examples siEpsin, siNECAP1 and siITSN1 were highlighted by colored lines to illustrate their differential effects on early stages v.s. *TfReff*. The data show that *CCP rate* was most affected by EAP KDs (reduced on average by 30%); whereas *TfRint* was only reduced by ~10%. *TfReff* was more sensitive than *TfRint*, but was still reduced on average by only <20%. The three highlighted EAPs, epsin1, NECAP1 and ITSN1, underline the distinguishing power of DASC vs biochemical CME measurements. While their individual KD resulted in an ~30% decrease in *CCP rate*, typical among the whole collection of KDs, they had differential effects on *TfReff*. Whereas KD of NECAP1 correspondingly decreased *TfReff* by ~ 24%, KD of ITSN1 and epsin1 caused only a minor decrease or insignificant change in *TfReff*. These examples indicated that early phenotypes can easily be obscured in non-stage-specific biochemical measurements.

We further illustrated this point for the whole collection of EAP KDs. For better visualization, we reduced the dimensionality of the extracted features using a principal component analysis (PCA) (implemented in Matlab’s function *pca*). The original data (Fig. 5A) contained 11 observations (11 EAP KDs) of 6 variables/dimensions (6 relative changes). First, the original observations were recentered, rescaled and projected into a new 2 dimensional PCA space, spanned by Component 1 and Component 2, which are linear combinations of the original 6 dimensions (Fig. 5C, implemented in Matlab’s function *biplot*). The variance of the original data was largely (>85%) maintained in this new space, shown by Component 1 (65.14% of total variance) and Component 2 (20.65% of total variance). Hence, the dimensionality reduction to a 2D space caused no substantial information loss from the original data. We then projected the original variables or dimensions into the two-component PCA space (Fig. 5C) and observed that Δ_r_*CCP rate* was almost perpendicular to Δ_r_*TfReff*. This striking orthogonality indicates manipulations of early CME stages have almost no effect on the bulk efficiency of CME. We conclude from this that defects in the CCP initiation and stabilization steps are compensated through redundant mechanisms that replenish transferrin receptor uptake. We supplemented the PCA with a correlation map (Fig. S5). Indeed, Δ_r_*CCP rate* among other early variables shows little correlation to Δ_r_*TfReff*. These comparisons highlight the value of DASC for increased sensitivity and greater phenotypic resolution over bulk biochemical measurements of cargo uptake, which can often obscure effects of EAP KD due to the resilience of CME.

## Discussion

Live cell imaging has revealed remarkable heterogeneity in the intensities and lifetimes of eGFP-CLCa-labeled CCPs in vertebrate cells, even amongst productive pits (13). Consequently, it has been challenging based on these parameters to comprehensively and objectively distinguish abortive coats (ACs) from *bona fide* CCPs, and thus to use them to define the roles of many uncharacterized EAPs in the dynamics of CCV formation. Here, we introduce DAS as a new feature space for describing CS dynamic behaviors, in which ACs and *bona fide* CCPs are accurately resolved. The DAS features exploit fluctuations in the inherently noisy intensity traces of individual CSs. The associated software pipeline, DASC, reliably separates dynamically, structurally and functionally distinct abortive and productive subpopulations without imposing any prior assumptions or the need for additional markers. While demonstrated on the classification of CSs during CME, the DASC framework is derived from first principles of thermodynamics describing entropy production during the assembly of macromolecular structures. Therefore, our tool will be applicable to any assembly process for which the addition and exchange of subunits can be traced.

Application of DASC to analyze the effects of knockdown of eleven early-acting endocytic accessory proteins identified diverse and significant phenotypes in discrete stage(s) of CCP progression, orthogonal to changes in conventionally used cargo uptake assays. Our findings establish the necessity of DASC for mechanistically dissecting early stages in CME dynamics and to study the numerous, as yet functionally undefined, endocytic accessory factors.

Characterization of the DASC-resolved AC and CCP subpopulations shows that ACs: i) have much lower average intensities than CCPs, ii) have much shorter average lifetimes than CCPs, iii) exhibit unregulated exponentially decaying lifetime distributions, as compared to the near-Rayleigh distributed CCP lifetimes, iv) contain fewer AP2 complexes than CCPs, v) recruit both clathrin and AP2 at a much slower rate than CCPs, and vi) acquire less curvature than CCPs. All of these features reproduce the properties of abortive coats inferred from previous studies (10, 16), thus both validating the robustness of DASC for distinguishing ACs from *bona fide* CCPs and providing unambiguous mechanistic insight into the factors required to stabilize nascent CCPs. Importantly, however, the distributions of each of these distinguishing properties have strong overlap between ACs and CCPs, preventing the use of any single or combined feature set as a marker for AC vs CCP classification. DASC is the only tool so far that can serve the purpose of stratifying individual CSs into these groups.

By applying this classification power to analyze early acting EAPs, we could assign their differential functions to specific stages of CCV formation even when single isoforms were individually depleted and bulk rates of cargo uptake were not or only mildly affected. Thus, DASC enables phenotypic assignment of individual EAPs to discrete stages of CME, but also reveals the existence of compensatory mechanisms (10, 44) and/or molecular redundancy (1) able to restore or maintain efficient cargo uptake. The resilience of CME to the effects of KD of individual components of the endocytic machinery is evident in the inability of multiple genome-wide screens based on ligand internalization assays to detect EAPs (45–48). Previous studies have shown that one compensatory mechanism triggered in cells expressing a truncated α-subunit lacking the EAP-binding appendage domain involves the isoform-specific activation of dynamin-1 (49). Thus, DASC will be a critical tool for future studies aimed at identifying other possible compensatory mechanisms able to restore transferrin receptor internalization.

We report a strong effect on the efficiency of *TfR* uptake in cells depleted of CALM and SNX9, whereas others have reported only minor or no effects (6–9). These differences may reflect cell type specific expression levels and/or activities of functionally redundant isoforms such as AP180 or SNX18 in the case of CALM and SNX9, respectively (8). Moreover, while not relevant to the work cited above, transferrin uptake assays that only measure the intracellularly accumulated ligand (e.g. *TfRint*) without taking into consideration changes in levels of surface receptor, as is frequently the case for FACS- or fluorescence microscopy-based assays, could miss or mis-interpret phenotypes (see Fig. S4). Importantly, the sensitivity of DASC to changes in early stages of CCP initiation and stabilization, enables detection of phenotypes even when single isoforms are depleted.

In summary, DASC classifies the previously unresolvable ACs and CCPs using data derived from single channel live cell TIRF imaging, thus providing an accurate measure of progression of CME through its early stages. This comprehensive and unbiased tool enables the determination of the distinct contributions of early EAPs to clathrin recruitment and/or stabilization of nascent CCPs. The stage-specific analysis by DASC is essential to characterize the functions of EAPs that were previously masked by detection limits and incompleteness of current experimental approaches. Going forward, DASC will be essential to functionally and comprehensively characterize the roles of the complete set of >70 EAPs in CME dynamics.

## Materials and Methods

### Computational flow of DAS analysis

1. Acquire intensity traces using cmeAnalysis (10) to analyze live-cell imaging movies. From the software output, determine the total number of traces, *N_tot_*, which includes both valid traces (*N* entries, *i.e*. always diffraction-limited with no consecutive gaps) and invalid traces (*N_iv_* entries, *i.e*. not always diffraction limited, and/or contain consecutive gaps) and calculate the *CS initiation rate (CS init.)*, which equals to *N_tot_*/(*A* · *T*), where *A* is the cell area and *T* = 451*s* is the duration of each movie. Repeat this step for control and all the experimental conditions that have been collected on the same day. It is critical that a new control be performed with each data set.
2. Include only ‘valid’ traces in the following DAS analysis (described below) to identify subpopulations of CSs.
3. Align each trace to its first frame, which is the first statistically significant detection (10). Then, for each trace, every intensity value is rounded to its nearest integer, *i* ∈ [1, *i_max_*](*a*. *u*.), where *i_max_* is the maximal rounded intensity among all the traces acquired on the same day.
4. Calculate conditional probabilities *W_t_*(*i*^⊖^|*i*) (*i.e*. increase in intensity from *t* to *t*+1) and *W_t_*(*i*|*i*^⊖^) (*i.e*.decrease in intensity from *t* to *t*+1), *t* ∈ [1, *T*], using the entire population of traces from the control condition:

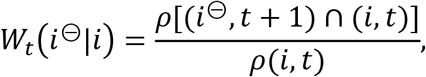

where *ρ*(*i*, *t*) is the probability of traces that reach (*i*, *t*), and *ρ*[(*i*^⊖^, *t* + 1) ∩ (*i*, *t*)] is the joint probability of traces that reach (*i*, *t*) but also reach (*i*^⊖^, *t* + 1). Conversely,

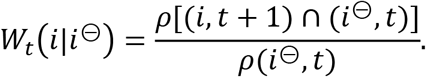 Note that large numbers of traces (>100,000), typically obtained from >10 movies per condition, are required to obtain stable values of *W_t_*.
5. Calculate the function *D*(*i*, *t*), based on eq. (2) (see main text). Note that the *D* function is only calculated once using control traces. The same *D*, which in essence serves as a ‘standard function’, will be applied to directly compare data between different conditions, if collected on the same day.
6. Convert each trace to a *D* series by substituting its intensity at each time frame (*i.e*. eq. (1)) into its *D* function (*i.e*. eq. (3)). Repeat this step for all conditions.
7. Calculate the three features *d*_1_, *d*_2_ and *d*_3_ of every *D* series, resulting in a *N* by 3 data set, where *N* is the total number of *D* series. Repeat this step for all the conditions.
8. Make the three features numerically comparable by normalizing *d*_1_, *d*_2_ and *d*_3_ from different conditions using means and standard deviations of the control. For any given condition, the normalized *d* reads:

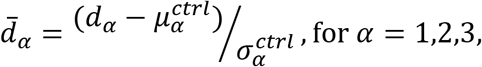

where 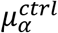 is the mean of all *d_α_* and 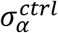 is the standard deviation of all *d_α_* in control condition. Apply the k-medoid method, using 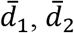 and 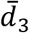 as features, to separate the traces from a single condition into *3* clusters, CCP, AC and OT, using Euclidean distance. k-medoid (implemented in Matlab’s function *kmedoids*) is chosen for its robustness over k-means. Repeat this step for all the conditions from the same day.
9. Calculate metrics such as lifetime and maximal intensity distributions and medians, population size, etc. for all traces within the same cluster. See more details of these calculations in the following sections. Repeat this step for all the conditions.
10. Calculate the fraction of CCPs, *CCP*% (the efficiency of CCP stabilization), as the population of CCPs, *n_CCP_*, divided by the entire population of valid traces, *N* = *n_CCP_* + *n_AC_* + *n_OT_*. Box plots with p-values are shown for CS init. and CCP% using Matlab’s exchange file function *raacampbell/sigstar* by Rob Campbell. Repeat this step for all the conditions.
11. Calculate *CCP rate* that equals to *n_CCP_*/(*A* · *T*) as the evaluation of the combined result of initiation and stabilization.
12. Evaluate statistical significance using Wilcoxon rank sum test (implemented in Matlab’s function *ranksum*).

### Statistical confidence bands of probability density functions based on bootstrapping

A new statistical analysis evaluating the variation of probability density function (pdf) is developed for the data in this paper, where movie-movie variation is considered to be the dominant source of variation. First, for a given choice of variable *x*, e.g. lifetime or maximal intensity in either CCP or AC subpopulations, *x* values pooled from all *N_m_* movies in a certain experimental condition are obtained. To equalize the contribution from different movies, *x* values in each movie are resampled to match the same size (*n_x_*) before pooling, where *n_x_* is the median of the *N_m_* movies’ CCP or AC number per movie. The pdf *p*(*x*) is then computed using Matlab’s function *ksdensity* (default kernel smoothing factor is applied to all pdf calculations). Next, to evaluate the movie-movie variation, the *N_m_* movies are bootstrapped to obtain *N_m_* resampled movies. *x* values from these bootstrapped movies are pooled to compute the first bootstrapped pdf 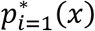 using *ksdensity*, where *i* indicates bootstrap number. Repeating this part 400 times, 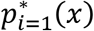 for *i* = 1… 400 are obtained. Finally, at any given value *x*, the 95% confidence band is obtained as a lower and upper bound [*p*_↓_(*x*), *p*^↑^(*x*)], where *p*_↓_ = 2.5^th^ percentile and *p*_↓_ = 97.5^th^ percentile of the 400 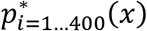 values. The final presentation of pdf is therefore *p*(*x*) as the main curve with the confidence band defined by *p*_↓_(*x*) and *p*^↑^(*x*).

### Normalized two-dimensional distributions

DAS plots (e.g. Fig. 2D), calculated as 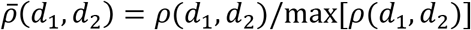 represent the 2D probability density normalized by maximum, where *ρ*(*d*_1_, *d*_2_) is the probability density in *d*_1_-*d*_2_ space, binned by Δ*d*_1_ = 0.2 and Δ*d*_2_ = 0.5. The normalized probability density projections of the data in the (*d*_1_, *d*_2_ *d*_3_) space in Fig. 2A is computed in the same way, adding bins of Δ*d*_3_ = 0.5.

The DAS difference maps (e.g. Fig. 3B) show the difference between the normalized 2D densities of two given conditions divided by their integrations (condition 1 as control),

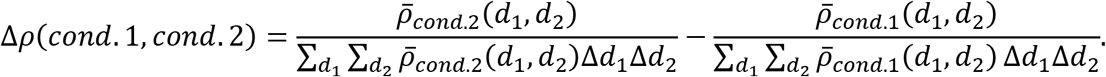

### Averaged intensity and Δz time course

For a given cohort lifetime *τ*, the traces within lifetime range *τ* ± 5*s* are averaged using the cohort method described in (10). The average values are presented as lines, and their error (standard deviation) as bands.

Using the microscopy setup illustrated in Fig. S3A, Epi and TIRF intensities over the lifetimes of each cohort (Fig. S3B-D) and errors of EPI and TIRF channels are obtained, *i.e*. *I_E_*(*t*) ± Δ*I_E_*(*t*) and *I_T_*(*t*) ± Δ*I_T_*(*t*). Following the approach developed by Saffarian and Kirchhausen (50), we then derived the distance between the center of the CS (*) and the initial position of assembled clathrin (+) as the invagination depth Δ*z* (Fig. S3A). For each cohort we calculated 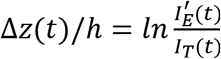, where the normalization factor is the characteristic depth of the TIRF field, *h* = 115*nm* based on our TIRF setting, similar to (26). 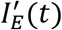 defines the Epi intensity trace adjusted to match the initial growth rate of clathrin measured in the TIRF intensity trace.

*I_E_* and *I_T_* are different in linear range of intensity measurement, *i.e*. the same intensity signal may have different readings from EPI and TIRF channel. To correct for this, *I_E_*(*t*) is adjusted along following protocol: 1) the data between the 2^nd^ and 10^th^ element in *I_E_*(*t*) and *I_T_*(*t*) are fitted by a 3^rd^ order polynomial *P_E_*(*t* = 2… 10s) and *P_T_*(*t* = 2… 10s) respectively. Then the initial growth rate for both channels is approximated as

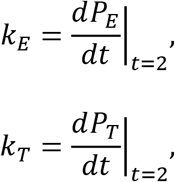

and *I_E_*(*t*) adjusted such that the growth rate of the corrected series *I*′_*E*_(*t*) matches *k_T_*, i.e., 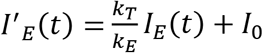 (*S*1, and *I*_0_ is an additive correction factor (see below). The averaged invagination depth is then extracted from the relation

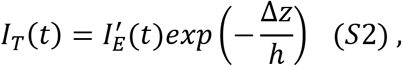

i.e.,

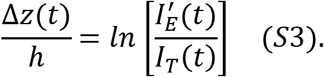

Considering the approximation that Δ*z*(*t* = 2) ≈ 0, *I*_0_ is obtained by substituting eq. (S1) into eq. (S3), and then replacing *I_E_* and *I_T_* at *t* = 2*s* with the corresponding fitted values from *P_E_* and *P_T_*,

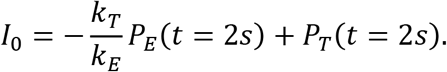

Δ*z*(*t*) is then expressed as a function of *I_T_* and the original *I_E_* with calculated parameter values,

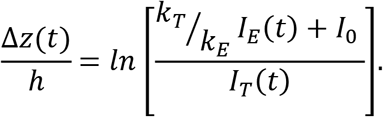

The error of Δ*z*(*t*) is obtained through error propagation for the two variables *I_E_*(*t*) ± Δ*I_E_*(*t*) and *I_T_*(*t*) ± Δ*I_T_*(*t*) using Matlab’s exchange file function *PropError* by Brad Ridder. Note that at early and late time points, high background but weak foreground intensity prohibits accurate calculation of *I_E_* and hence Δ*z*(Fig. 3F and S3). We also detected too few ACs in the 40s cohort for robust analysis (Fig. S3).

### Cell culture and cell engineering

ARPE19 and ARPE-19/HPV-16 (ATCC^®^ CRL-2502™) cells were obtained from ATCC and cultured in DMEM/F12 medium with 10% (v/v) FBS at 37°C under 5% CO2. ARPE-19/HPV-16 cells were infected with recombinant lentiviruses encoding eGFP-CLCa in a pMIEG3 vector, and sorted by FACS after 72 hours (10). AP2 reconstitution was achieved by infecting the eGFP CLCa-expressed ARPE-19/HPV-16 cells (ARPE_HPV16 eGFP_CLCa) with retroviruses encoding siRNA resistant WT or PIP2- (K57E/Y58E) AP2 alpha subunit in a pMIEG3-mTagBFP vector and FACS sorted based on BFP intensity (17). CALM reconstitution was achieved by infecting ARPE-19/HPV-16 eGFP-CLCa cells with retroviruses encoding siRNA resistant WT CALM in a pBMN vectors gifted from Dr. David Owen (21) (Cambridge, UK) and selected in 0.25 mg/ml hygromycin B for a week. Western blotting was used to confirm reconstituted-protein expression and knockdown efficiency of the generated cell lines using anti-alpha-adaptin (Thermo Fisher Scientific, #AC1-M11) and anti-CALM (Abcam, #ab172962) antibodies. APRE19 cells with stable expression of mRuby2-CLCa and α-eGFP-AP2 were also generated via lenti- and retroviral transduction, respectively.

### siRNA transfection

200,000 ARPE-19/HPV-16 cells were plated on each well of a 6-well plate for ≥ 3 hours before transfection. Transfections for siRNA knockdown were assisted with Lipofectamin RNAiMAX (Life Technologies, Carlsbad, CA). Briefly, 6.5 μl of Lipofectamin RNAiMax and 5.5 μl of 20 μM siRNA were added separately into 100 μl OptiMEM and incubated separately for 5 min at room temperature. SiRNA were next mixed with lipofectamin RNAiMAX and incubated at room temperature for another 10 min before being added dropwise to the cells with fresh medium. Measurements were performed at day 5 after plating cells following two rounds of siRNA transfection (time gap = 24-48 hrs between transfections).

Western blotting confirmed that the knockdown efficiency for all proteins was over 80%. Control cells were transfected in parallel with control siRNA (siCtrl) purchased from QIAGEN (Germantown, MD).

### Transferrin receptor internalization assay

Internalization of transferrin receptor was quantified by in-cell ELISA following established protocol (39). ARPE-19/HPV-16 cells were plated in 96 well plates (15,000 cells/well, Costar) and grown overnight. Before assay, cells were starved in PBS4+ (1X PBS buffer with addition of 0.2% bovine serum albumin, 1mM CaCl2, 1mM MgCl2, and 5mM D-glucose) for 30min at 37°C incubator with 5% CO2 and then cooled down to 4°C and supplied with 100μl 5μg/ml HTR-D65 (anti-TfR mAb) (51). Some cells were kept at 4°C for the measurement of surface-bound HTR-D65, while some cells were moved to 37°C water bath for 10min internalization and then acid washed to remove surface-bound HTR-D65. All cells were fixed with 4% paraformaldehyde (PFA) (Electron Microscopy Sciences, PA) and penetrated with 0.1% Triton-X100 (Sigma-Aldrich). After blocking with Q-PBS (PBS, 2% BSA, 0.1% lysine, 0.01% saponin, pH 7.4) for 30min, surface and internalized HTR-D65 was probed by HRP-conjugated goat-anti-mouse antibody (Sigma-Aldrich). Color developed after adding OPD solution (Sigma-Aldrich) and absorbance was read at 490nm (Biotek Synergy H1 Hybrid Reader).

Statistical significance of changes in internalized and surface-bound transferrin receptors (*TfRint* and *TfRsuf*) were obtained by two-sample t-test (implemented in Matlab’s function *test2*). Statistical significance of changes and 95% confidence intervals in efficiency of transferrin receptor uptake (*TfReff* = *TfRint/TfRsuf*) were obtained using a statistical test for ratios (52) (implemented in a customized Matlab’s function).

### Microscopy imaging and quantification

Total Internal Reflection Fluorescence (TIRF) Microscopy imaging was conducted as previously described (16). Cells were grown on a gelatin-coated 22×22mm glass (Corning, #2850-22) overnight and then mounted to a 25×75mm cover slide (Thermo Scientific, #3050). Imaging was conducted with a 60X, 1.49-NA Apo TIRF objective (Nikon) mounted on a Ti-Eclipse inverted microscope equipped with an additional 1.8X tube lens, yielding a final magnification of 108X. Perfect focus was applied during time-lapsed imaging. For epi/TIRF imaging, nearly simultaneous two channel (488 epifluorescence/TIRF) movies were acquired with multi-dimension acquisition (MDA). Movies were acquired at the rate of 1 frame/s. cmeAnalysis was applied for CCP detection and tracking (10, 23, 26).

## Acknowledgements

We thank Rosa Mino for generating the α-eGFP-AP2/mRuby-CLC expressing RPE cells and Schmid lab members for helpful discussion. This work was supported by NIH grants GM73165 to SLS and GD, MH61345 to SLS and GM067230 to GD. ZC was supported by Welch grant I-1823 to SLS.

## Supplemental figures

**Figure S1.**
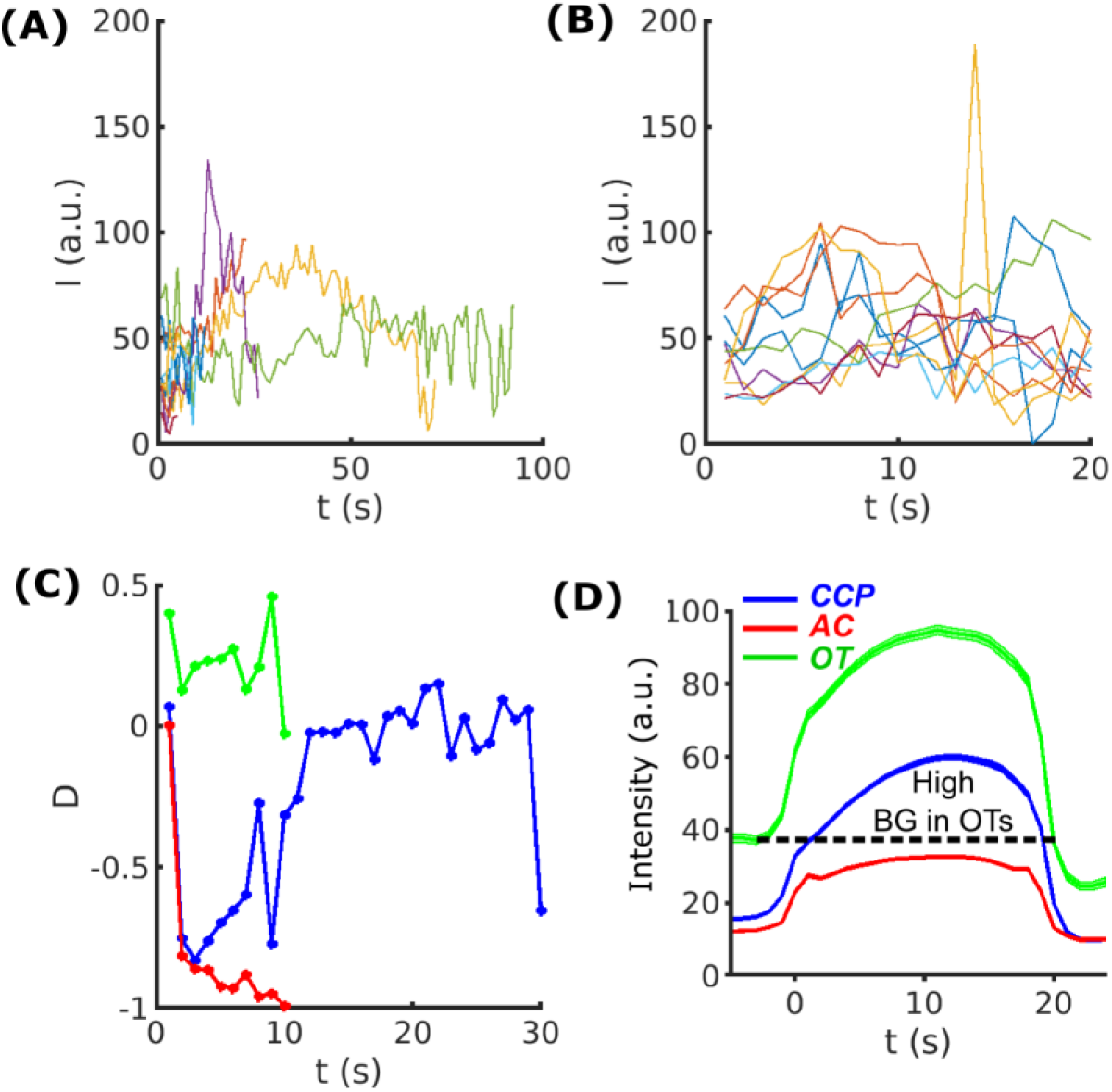
(A) Ten randomly selected traces of eGFP-CLCa intensity at CSs in WT cells. (B) Ten randomly selected traces with the same lifetime *τ* = 20 seconds from the same cells as in (A). (C) *D* values as time series read from the color map corresponding to the three traces in Fig. 1C in the main text. Color scheme: CCP (blue), AC (red) and OT (green). (D) 20s cohort of CCP, AC and OTs. Same color scheme as in (C). High background (BG) in dashed line observed in OTs.

**Figure S2.**
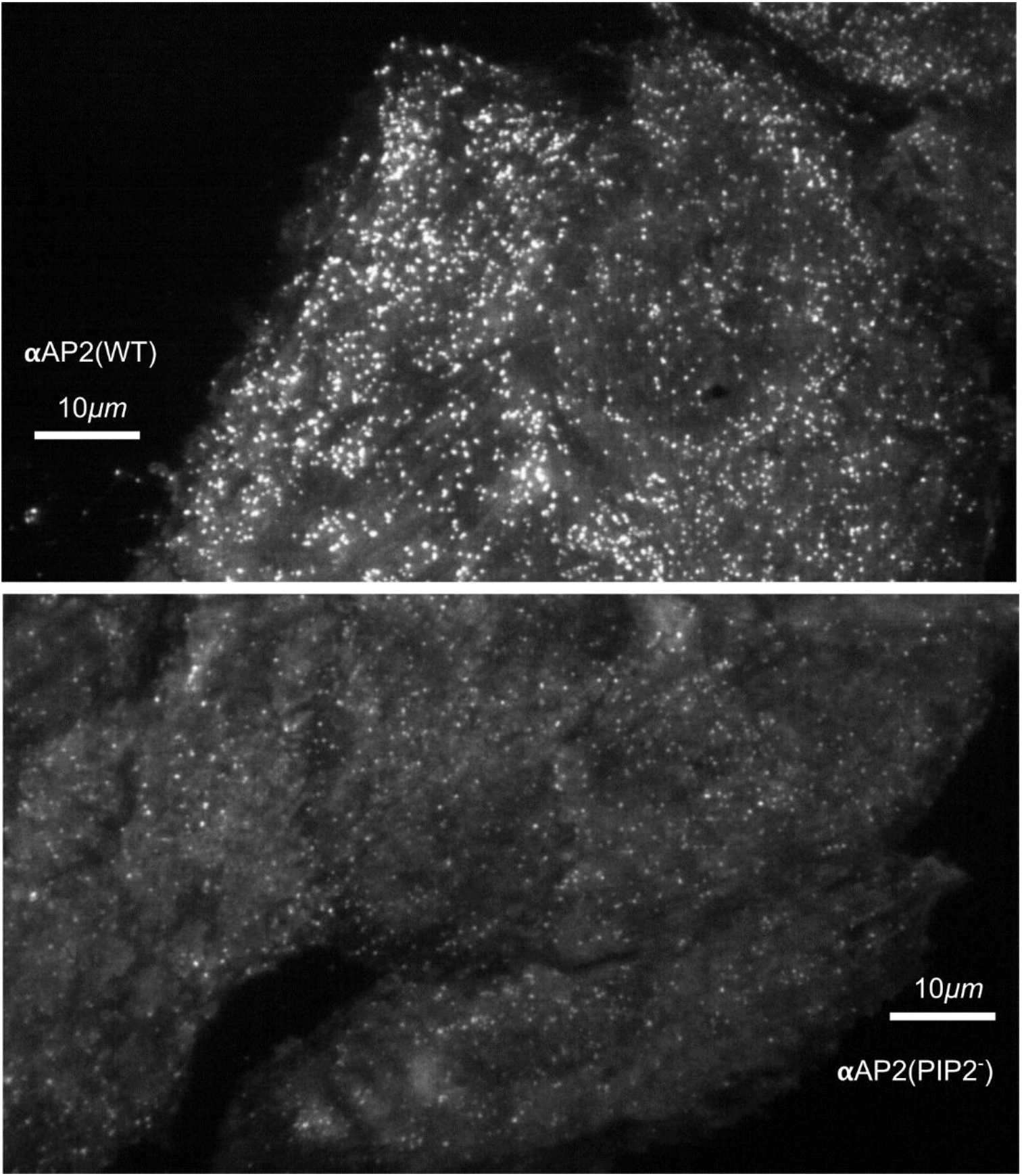
Single frame from movies of αAP2(WT) and αAP2(PIP2^-^) cells. Note that CCPs in the αAP2(PIP2^-^) cells are much dimmer potentially altering the ability to detect valid CS initiation events.

**Figure S3.**
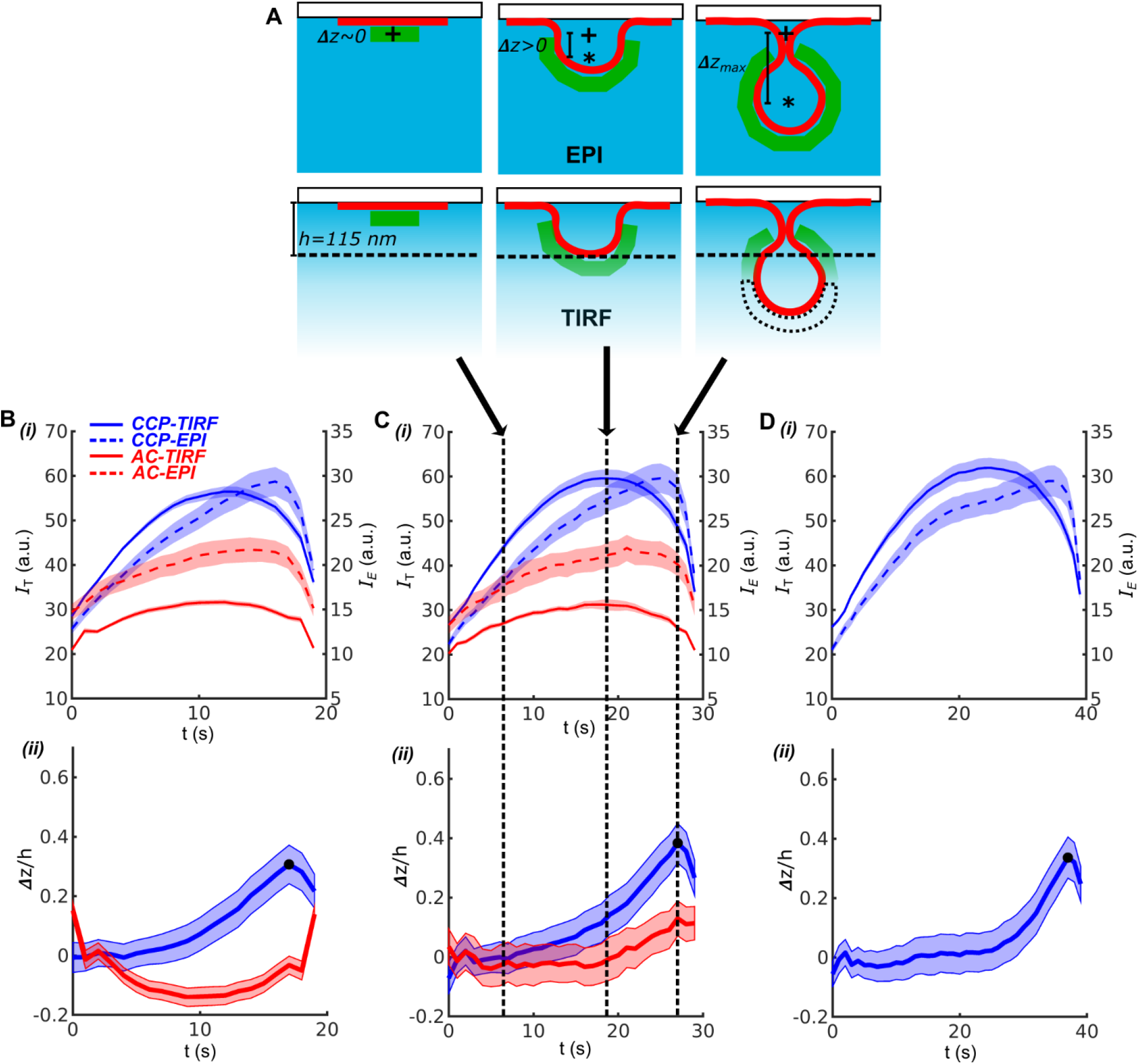
DASC combined with EPI-TIRF approach reveals CME invagination kinetics. (A) Schematic of CCP in EPI and TIRF microscopy at 0, intermediate and maximal invagination depth (Δ*z*). ‘+’ as starting point and ‘*’ as center of mass of CCP during invagination. TIRF characteristic depth *h* = 115*nm*. (B) i. 20s cohorts of CCPs in TIRF channel (blue, solid line) and EPI (blue, dashed), and ACs in TIRF (red, solid) and EPI (red, dashed); ii. Δ*z*(*t*)/*h* time course of CCPs (blue) and ACs (red) derived from EPI-TIRF cohorts in i, Δ*z_max_* indicated as dark dot. (C) i-ii Same as (B) for 30s cohorts and *Δz(t)/h*. (D) i-ii Same as (B) for 40s cohorts and Δ*z*(*t*)/*h* but without ACs. Shaded area as 95% confidence interval (Materials and Methods).

**Figure S4.**
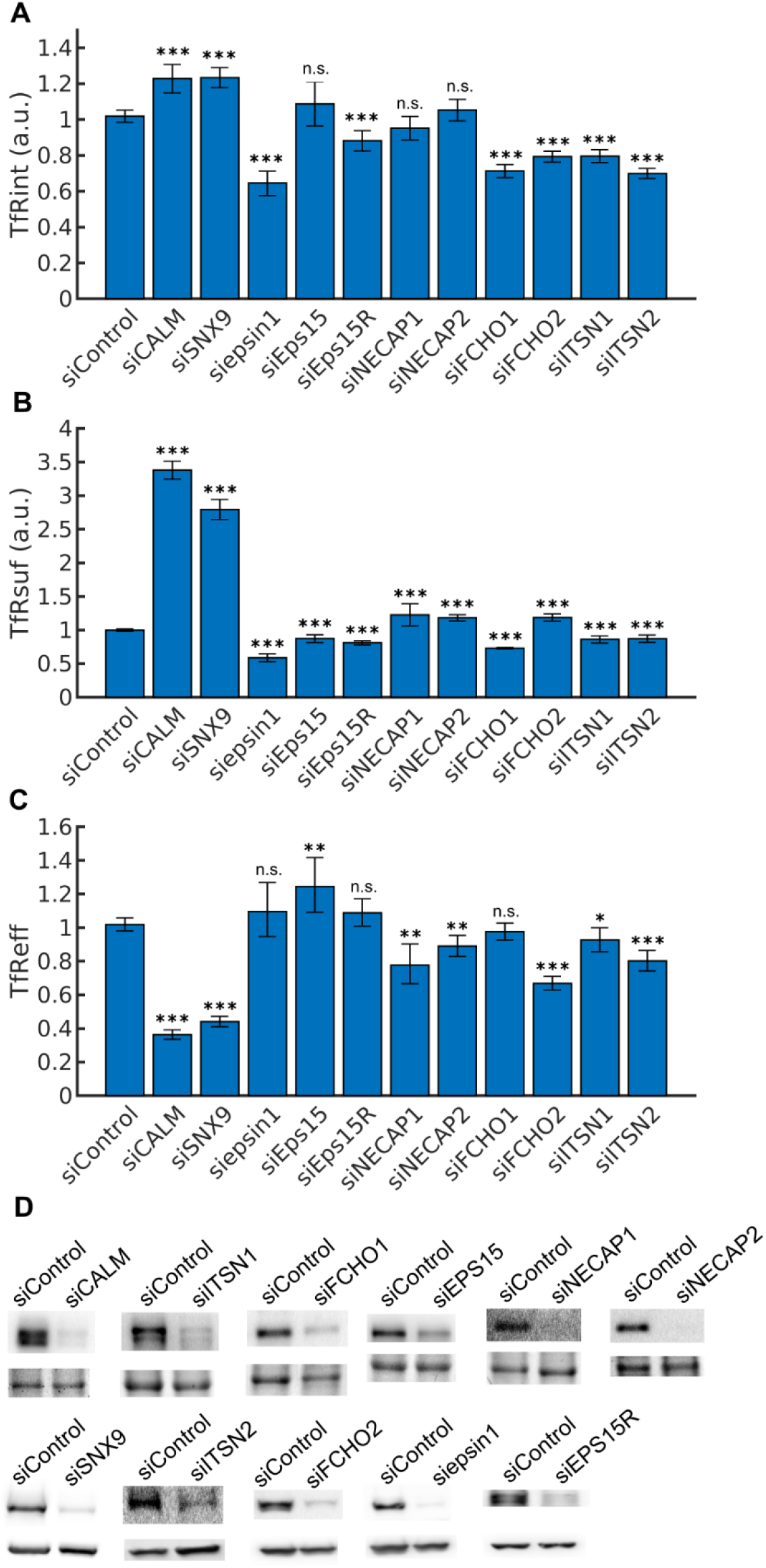
Measurements of transferrin receptor uptake and siRNA knockdown efficiency. (A) Internalized transferrin receptors (*TfRint*) after 10 min in arbitrary unit (a.u.). (B) Surface bound transferrin receptors (*TfRsuf*) (a.u.). (C) Efficiency of transferrin receptor uptake (*TfReff*) as ratio of *TfRint* over *TfRsuf*. Error bars represent 95% confidence intervals. Statistical significance of *TfRint* and *TfRsuf* are obtained using 2-sample t-test. A statistical test for ratios is applied to calculate the significance of *TfReff*, see Materials and Methods. (D) KD efficiency of 11 EAPs shown by western blots.

**Figure S5.**
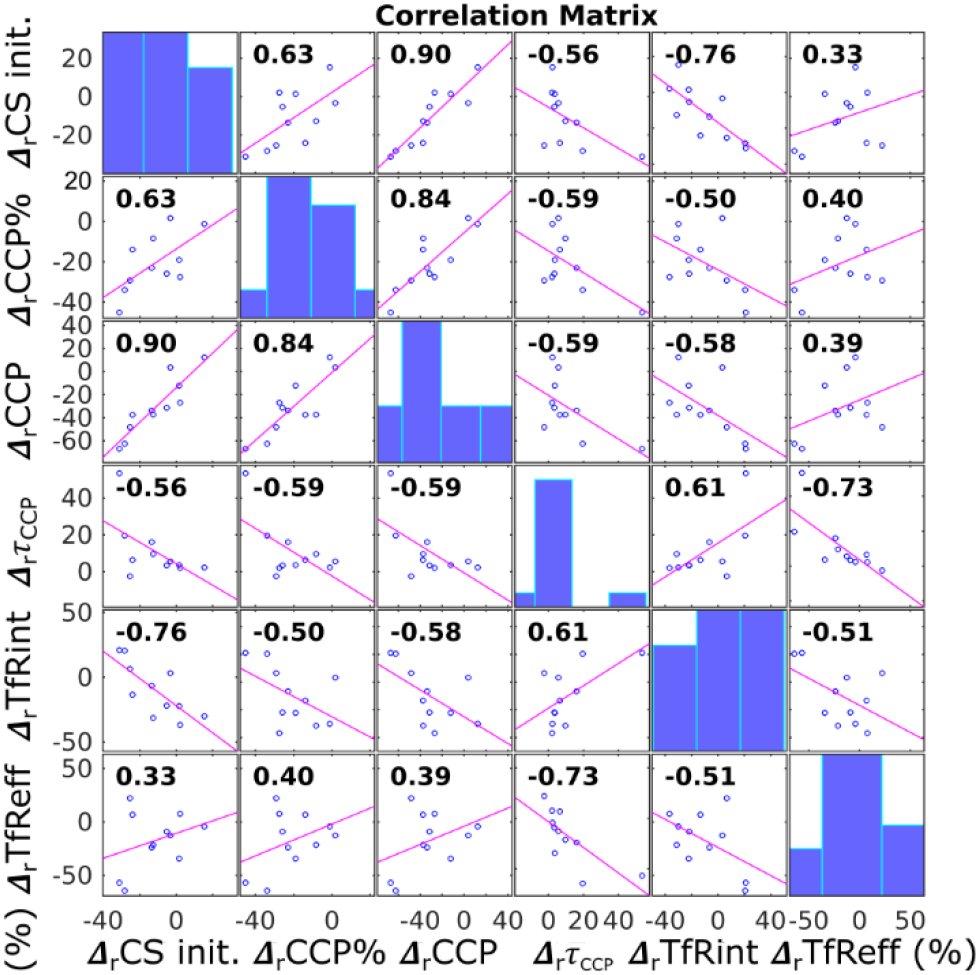
Correlation matrix of 6 variables in Fig. 5A. Diagonal bar graphs showed histogram of individual variable values. Off-diagonal graphs showed pair-wise Pearson linear correlation coefficient. Implemented in Matlab’s function *corrplot*.

